# The endoplasmic reticulum-resident protein TMEM-120/TMEM120A promotes fat storage in *C. elegans* and mammalian cells

**DOI:** 10.1101/2021.06.29.450322

**Authors:** Yan Li, Siwei Huang, Xuesong Li, Xingyu Yang, Ningyi Xu, Jianan Qu, Ho Yi Mak

**Affiliations:** Division of Life Science, The Hong Kong University of Science and Technology, Hong Kong SAR, China; Biophotonics Research Laboratory, Department of Electronic and Computer Engineering, The Hong Kong University of Science and Technology, Hong Kong SAR, China; The MOE Key Laboratory of Biosystems Homeostasis & Protection and Innovation Center for Cell Signaling Network, Life Sciences Institute, Zhejiang University, Hangzhou, Zhejiang, China

**Keywords:** TMEM-120, TMEM120A, endoplasmic reticulum, lipid droplets

## Abstract

The synthesis of triacylglycerol (TAG) is essential for the storage of excess fatty acids, which can subsequently be used for energy or cell growth. A series of enzymes act in the endoplasmic reticulum (ER) to synthesize TAG, prior to its transfer to lipid droplets (LDs), which are conserved organelles for fat storage. Here, we report that the deficiency of TMEM-120/TMEM120A, a protein with 6-transmembrane helices, retards TAG synthesis and LD expansion in *C. elegans*. A missense mutation near the predicted coenzyme A binding site of TMEM-120 confers strong loss of function phenotypes. GFP fusion proteins of TMEM-120, expressed at the endogenous level in live worms, were observed throughout the ER network. Using Stimulated Raman Scattering, we discovered a specific requirement of TMEM-120 in the storage of exogenous fatty acids in LDs. Knockdown of TMEM120A impedes adipogenesis of pre-adipocytes in vitro, while its over-expression is sufficient to promote LD expansion. Pharmacological studies indicate that TMEM120A most likely acts upstream of diacylglycerol O-acyltransferase 1 (DGAT1). Our results suggest that TMEM-120/TMEM120A plays a conserved role in increasing the efficiency of TAG synthesis.

## Introduction

Excess fatty acids from de novo lipogenesis or the diet can be incorporated into neutral fat, such as triacylglycerol (TAG), via the glycerol-3-phosphate (G3P) or the monoacylglycerol (MAG) pathway (Yen et al., 2008). Common to both pathways is the addition of fatty acyl-Coenzyme-A (acyl-CoA) molecules to specific positions of the glycerol backbone, by acyltransferases (Coleman and Lee, 2004). Based on biochemical and imaging analyses, all TAG biosynthetic enzymes can be found in the endoplasmic reticulum (ER). Accordingly, newly synthesized TAG accumulates between ER membrane leaflets before the directional budding of the cytoplasmic leaflet to form nascent lipid droplets (LDs), which are conserved organelles for fat storage (Henne et al., 2018; Thiam and Ikonen, 2021; Walther et al., 2017). Additional TAG synthesis occurs at ER-LD contacts to support LD expansion (Cao and Mak, 2020; Olzmann and Carvalho, 2019; Schuldiner and Bohnert, 2017). Although the core ensemble of TAG biosynthetic enzymes have been well-defined, it is unknown if additional proteins are required for the maximal efficiency of TAG synthesis.

TMEM120A (alternatively known as NET29 or TACAN) is a member of a conserved family of transmembrane proteins that had been assigned seemingly unrelated functions. It was first reported in a proteomic study of the nuclear envelop (Schirmer et al., 2003). Subsequent functional analysis indicates that TMEM120A, and its paralog TMEM120B, is required for the differentiation of 3T3-L1 pre-adipocytes into mature adipocytes (Batrakou et al., 2015). More recently, TMEM120A was reported to act at the cell surface as an ion channel that is sensitive to mechanical cues (Beaulieu-Laroche et al., 2020). However, four independent studies did not support such conclusion (Ke et al., 2021; Niu et al., 2021; Rong et al., 2021; Xue et al., 2021). Instead, structural and biochemical analyses indicate that TMEM120A forms a symmetrical homodimer and each protomer binds specifically to a coenzyme-A (CoASH) molecule (Niu et al., 2021; Rong et al., 2021; Xue et al., 2021). These observations led to the proposal that TMEM120A has an undefined role in lipid metabolism, which correlates with its requirement for adipogenesis (Batrakou et al., 2015; Czapiewski et al., 2021).

We identified mutant worms that were deficient of the sole *C. elegans* ortholog of TMEM120A from an unbiased forward genetic screen. Here, we present evidence that *C. elegans* TMEM-120 acts at the ER to promote TAG synthesis.

## Results and Discussion

### TMEM-120 is required for TAG accumulation and LD expansion

We have previously shown that lipid droplets (LDs) undergo continuous expansion in *C. elegans daf-22*/thiolase mutant worms, owing to a block in the peroxisomal β-oxidation pathway (Zhang et al., 2010). These worms accumulate more triacylglycerol (TAG), which is synthesized by the concerted action of ACS-22/acyl-CoA synthetase and DGAT-2/diacylglycerol O-acyltransferase (Xu et al., 2012). Accordingly, loss of *acs-22* or *dgat-2* function attenuates LD expansion of *daf-22* mutant worms. In a genetic screen for additional *daf-22* suppressors, we identified a complementation group that consisted of two recessive alleles, *hj49* and *hj50*. Molecular cloning revealed lesions in a gene annotated as M01G5.3, hereafter named *tmem-120*, which encodes a predicted transmembrane protein that is homologous to mammalian TMEM120A and TMEM120B (Batrakou et al., 2015) (Fig. S1A). The *hj50* nonsense allele (Glutamine 290 to amber) caused ~90% reduction of *tmem-120* mRNA level (Fig. S1A-B), possibly due to nonsense mediated decay. Therefore, it is a strong loss of function allele. The *hj49* missense allele caused the substitution of a conserved Glycine to Glutamate (G195E) (Fig. S1A). Based on the recently determined structures of TMEM120A, this conserved Glycine residue locates in a flexible linker immediately N-terminal to a coenzyme-A (CoASH) binding site (Niu et al., 2021; Rong et al., 2021; Xue et al., 2021). Therefore, its replacement with Glutamate may alter the conformation of the linker and in turn affect the orientation of residues that constitute the CoASH binding site (including W193 of human TMEM120A and W197 of *C. elegans* TMEM-120). Since the phenotypes of mutant worms carrying *hj49* and *hj50* were indistinguishable (Fig. S1C-E), we concluded that the G195E substitution severely compromised TMEM-120 function. In subsequent experiments, we used worms carrying the *hj50* nonsense allele for phenotypic analysis.

To visualize LDs of wild type and mutant animals, we used a recently developed Stimulated Raman Scattering (SRS) microscopy system (Li et al., 2015). We focused on detecting C-H bond vibration from TAG, which was highly concentrated in LDs. In agreement with previous results based on the use of vital dye or fluorescent protein markers (Xu et al., 2012), LDs in *daf-22* mutant worms were larger than those in wild type worms (Fig. 1A). The loss of TMEM-120 function reduced LD size and blocked LD expansion in wild type and *daf-22* mutant worms (Fig. 1A and C). Similar results were observed when we used DHS-3::mRuby as a fluorescent LD marker (Zeng et al., 2020) (Fig. 1F). Next, we quantified SRS signals, which were proportional to TAG content (Wang et al., 2011). We detected 23% more SRS signals in *daf-22* mutant than wild type worms (Fig. 1B and D). In contrast, *tmem-120* and *daf-22*; *tmem-120* mutant worms have 26% and 14% less SRS signals than wild type worms, respectively (Fig. 1B and D). To complement our imaging approach, we used liquid chromatography-mass spectrometry (LC-MS) to determine that *tmem-120* mutant worms had almost 40% less TAG than wild type worms (Fig. 1E). Such reduction of TAG was unlikely due to an alteration of feeding rate (Fig. S1F). Taken together, our results indicate that TMEM-120 supports TAG accumulation and LD expansion.

**Figure 1.**
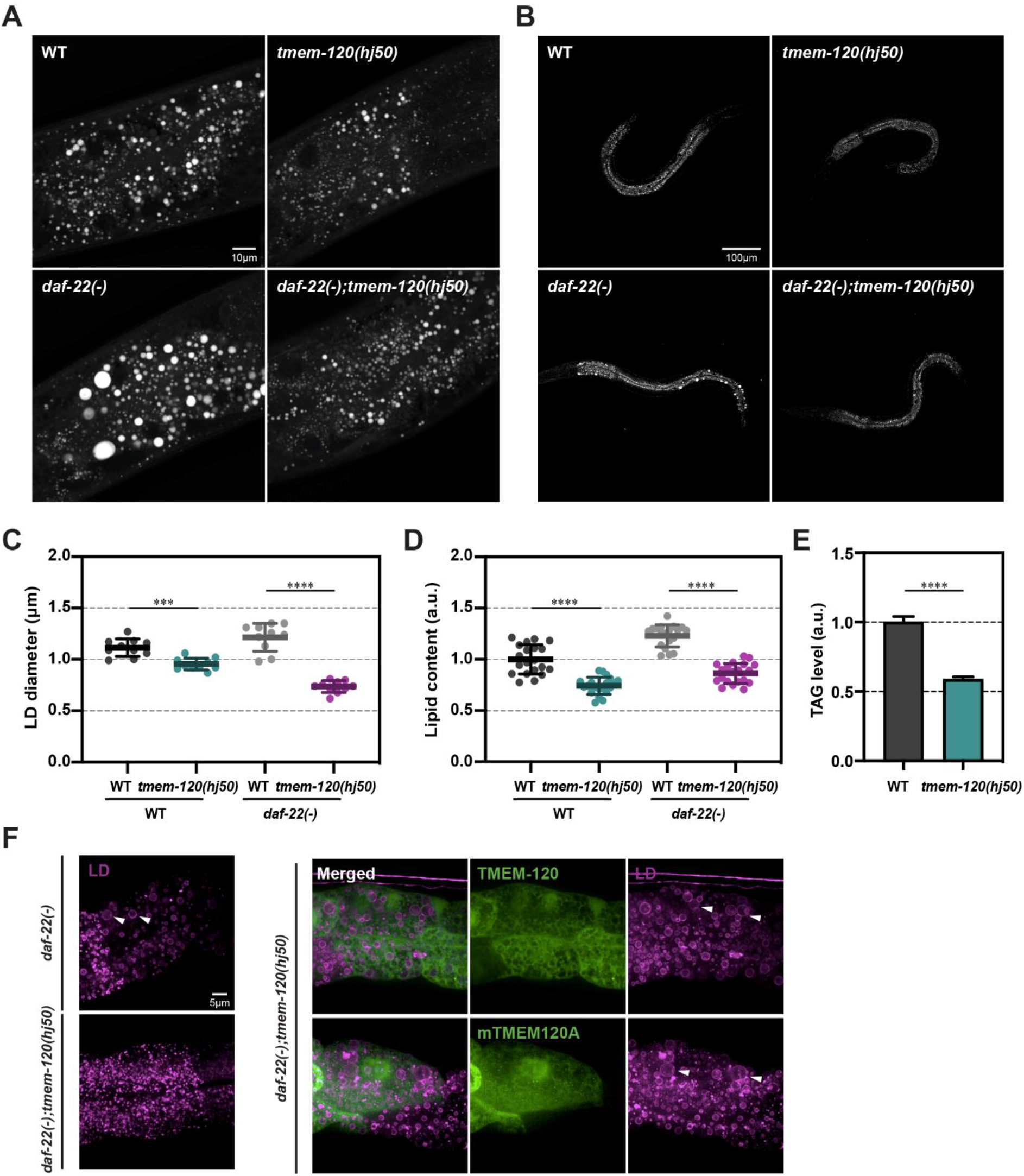
TMEM-120 promotes TAG synthesis and LD expansion. (A) Visualization of LDs in wild type (WT), *tmem-120*(*hj50*), *daf-22*(*ok693*) and *daf-22*(*ok693*); *tmem-120*(*hj50i*) young adult animals by Stimulated Raman Scattering (SRS). Representative images of a single focal plane of the first and second intestinal segments are shown. The anterior end of the worm is toward the left. For simplicity, *daf-22*(*ok693*) will be referred to as *daf-22*(-) thereafter. (B) As in (A), but with representative images of entire larval stage L4 worms at a focal plane with the strongest SRS intensity. (C) Quantification of LD diameter of WT, *tmem-120*(*hj50*), *daf-22*(-) and *daf-22(-); tmem-120(hj50)* larval L4 stage worms (n = 10 for each strain), using DHS-3::mRuby (*hj200*) as a LD marker. Each data point represents the average LD diameter of an individual worm. Total number of LDs quantified: WT = 2033, *tmem-120*(*hj50*) = 1561, *daf-22*(-) = 1331 and *tmem-120*(*hj50*); *daf-22*(-) = 1337. (D) Label free quantification of neutral lipid content by SRS in WT (n=20), *tmem-120*(*hj50*) (n=20), *daf-22*(-) (n=19) and *daf-22*(-); *tmem-120*(*hj50*) (n=20) worm shown in (B). Each data point represents neutral lipid content of an individual worm. The mean value of WT worms is assigned as 1. a.u. = arbitrary unit. (E) Quantification of TAG level in WT and *tmem-120*(*hj50*) L4 stage animals by LC-MS. Three independent biological samples for each group. Each group consists of ~2000 worms. The mean value of WT worms is assigned as 1. (F) Visualization of LDs in L4 stage worms using DHS-3::mRuby (*hj200*) as a LD marker. Representative images of *daf-22(-), daf-22(-); tmem-120*(*hj50*), *daf-22(-); tmem-120*(*hj50*); *Ex[vha-6p::tmem-120::sl2::gfp]* and *daf-22(-); tmem-120*(*hj50*); *Ex[vha-6p::mouse tmem120A::sl2::gfp]* are shown. The GFP is expressed from the same operon as TMEM-120 or mouse TMEM120A, but not as a fusion protein. Each representative image is a projection of 7.5μm z stack with the second intestinal segment in the center area. Enlarged LDs are indicated by white arrowheads. For all graphs, bars or horizontal lines represent mean ± SD. Statistical analysis: (C-D) two-way ANOVA followed by Sidak’s multiple comparisons test; (E) unpaired t test. ***p < 0.001; ****p < 0.0001.

Next, we sought to determine if mouse TMEM120A plays a conserved role in regulating fat storage, even though it was reported to be a mechanosensory channel (Beaulieu-Laroche et al., 2020). To this end, we expressed mouse TMEM120A in *daf-22*; *tmem-120* mutant worms. We found that TMEM120A could rescue the LD phenotype in a similar manner as *C. elegans* TMEM-120 (Fig. 1F). As a result, large LDs re-appeared in intestinal cells of *daf-22*; *tmem-120* mutant worms. Therefore, we conclude that TMEM-120 and TMEM120A share a deeply conserved function of regulating cellular fat storage.

### TMEM-120 promotes the incorporation of fatty acids into TAG

The cellular neutral lipid homeostasis is dependent on a balance between TAG synthesis and mobilization at LDs. The TAG synthesis in turn relies on the availability of dietary or de novo synthesized fatty acids. We sought to determine if the decrease in neutral lipid content was caused by a decrease in TAG accumulation or accelerated TAG mobilization by lipolysis in *tmem-120* mutant worms. To differentiate these possibilities, we used Stimulated Raman Scattering (SRS) to detect exogenously supplied deuterium-labeled fatty acids (Fu et al., 2014; Li et al., 2019). We tuned our system to detect C-D bond vibrations to avoid the interference from endogenous fatty acids. To measure TAG accumulation, we fed young adult wild type and mutant worms with deuterium-labeled monounsaturated oleic acid (OA-d_34_) and imaged live worms by SRS at regular intervals over a period of 30 hours (Fig. 2A). We detected progressive increase of SRS signals from LDs in both strains tested (Fig. 2C). However, the rate of increase in *tmem-120* worms was significantly slower than that in wild type worms (Fig. 2E). Similar observations were made when worms were fed deuterium-labeled saturated palmitic acid (PA-d_31_) (Fig. S2A, C and E). We conclude that loss of TMEM-120 function impairs the incorporation of exogenous fatty acids into TAG.

**Figure 2.**
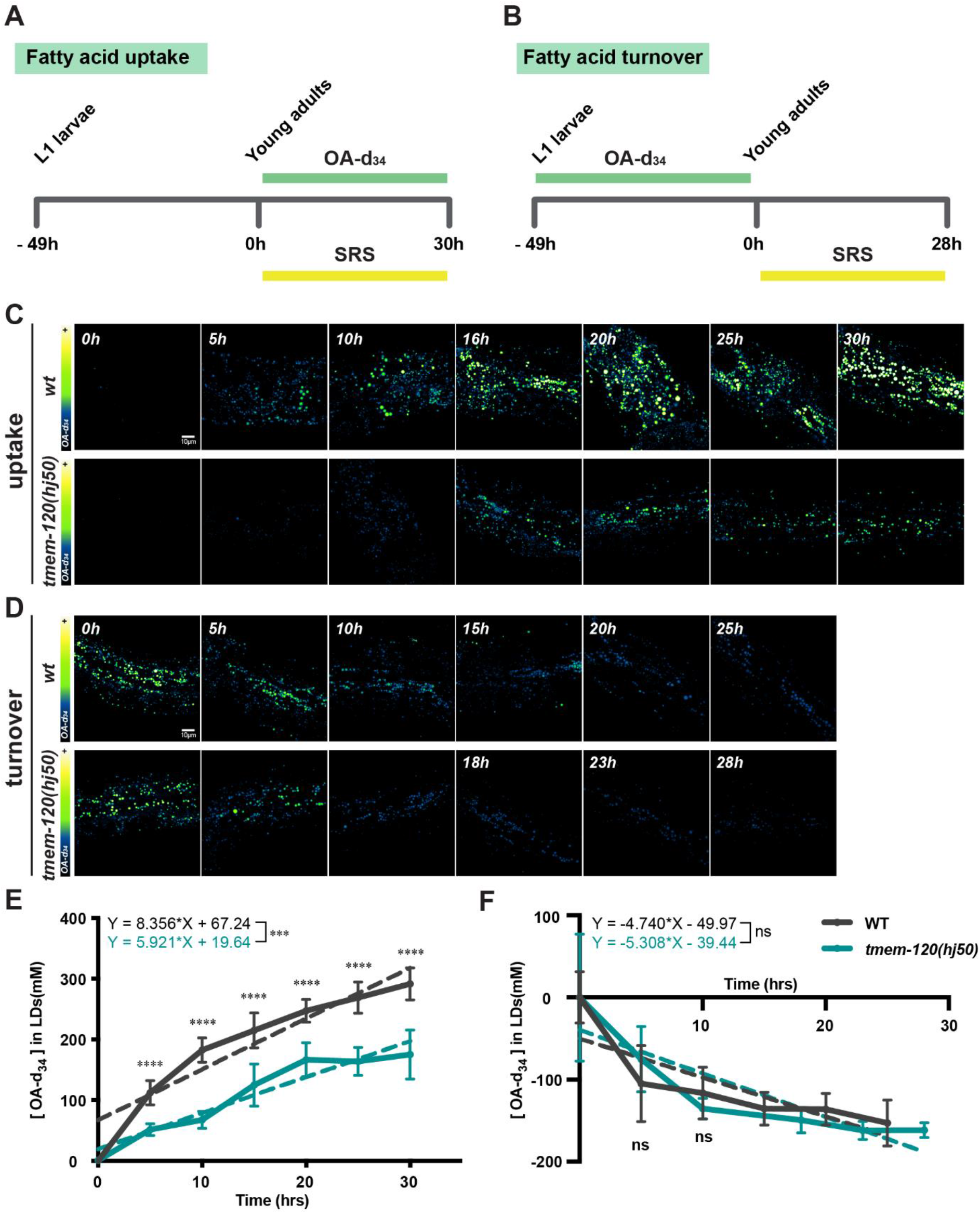
TMEM-120 promotes the incorporation of fatty acids into TAG. (A-B) The experimental design for monitoring deuterated oleic acid-d_34_ (OA-d_34_) uptake (A) or turnover (B). (C-D) Visualization of OA-d_34_ incorporation (C) or turnover (D) by SRS in wild type (wt) and *tmem-120*(*hj50*) worms. Representative images of a layer with the strongest SRS signal in the first and second intestinal segments are shown. (E) Quantification of OA-d_34_ uptake in WT and *tmem-120*(*hj50*) worms shown in (C). n = 5 to 9 for each group at each time point. Straight dashed lines and equations were generated based on linear regression analysis of each group. (F) Quantification of OA-d_34_ turnover in WT and *tmem-120*(*hj50*) worms shown in (D). n = 7 to 10 for each group at each time point. Statistical analysis: unpaired t test (for each time point). ns, not significant; ***p < 0.001.

To measure lipolysis, we fed newly hatched larval stage L1 worms with deuterium-labeled monounsaturated oleic acid (OA-d_34_) until they reached the young adult stage. We then removed labeled fatty acids from the diet of these animals and imaged them by SRS at regular intervals (Fig. 2B). The dissipation of SRS signals over time reflected the rate of lipolysis, as the labeled fatty acids stored as TAG in LDs were metabolized. We found that the rate of decrease of SRS signals was not significantly different between wild type and *tmem-120* mutant worms (Fig. 2D and F). Similar observations were made when worms were fed with saturated palmitic acid (PA-d_31_) (Fig. S2B, D and F). We conclude that lipolysis is not altered in *tmem-120* mutant worms. Taken together, our results indicate that the reduction of neutral lipid content in these animals was primarily due to the retardation of fatty acid incorporation into TAG.

### TMEM-120 is an ER resident protein

Based on TOPCONS analysis (Tsirigos et al., 2015), the *C. elegans* TMEM-120 and its human orthologs share the same predicted membrane topology, with their N- and C-termini facing the cytoplasm (Fig. 3A). According to its cryo-EM structure, membrane embedded TMEM120A has 6 transmembrane helices and forms homodimers (Ke et al., 2021; Niu et al., 2021; Rong et al., 2021; Xue et al., 2021). Intriguing, a consensus has yet to emerge regarding the subcellular localization of TMEM120A. Mammalian TMEM120A was previously reported to reside at the nuclear envelope (Batrakou et al., 2015) or the endoplasmic reticulum (ER) (Cho et al., 2020). In contrast, the assignment of TMEM120A as a mechanosensory channel placed it at the plasma membrane (Beaulieu-Laroche et al., 2020). We sought to determine the localization of *C. elegans* TMEM-120 when it was expressed at the endogenous level in live animals. To this end, we used CRISPR mediated genome editing (Arribere et al., 2014) to insert the coding sequence of the green fluorescent protein (GFP) into the endogenous *tmem-120* locus (Fig. 3B). We generated two independent knockin alleles, *hj239* and *hj270* (Fig. 3B). In one strain, GFP was fused to the N-terminus of TMEM-120 via a flexible linker. In a second strain, GFP was expressed as part of the predicted C-terminal cytoplasmic tail, leaving the extreme C-terminus unaltered. This design was necessitated by the presence of a KxHxx motif at the C-terminus of TMEM-120, which could function similarly as the KxKxx motif for ER retention (Jackson et al., 1990; Ma and Goldberg, 2013). Both TMEM-120 fusion proteins were functional because their expression did not significantly alter the lipid content of *daf-22* mutant worms, as observed when the function of TMEM-120 was lost (Fig. S3A-B). We used established markers of intestinal ER and LDs to ascertain the localization of TMEM-120 (Fig. 3C-D) and found that it co-localized extensively with the ER marker.

**Figure 3.**
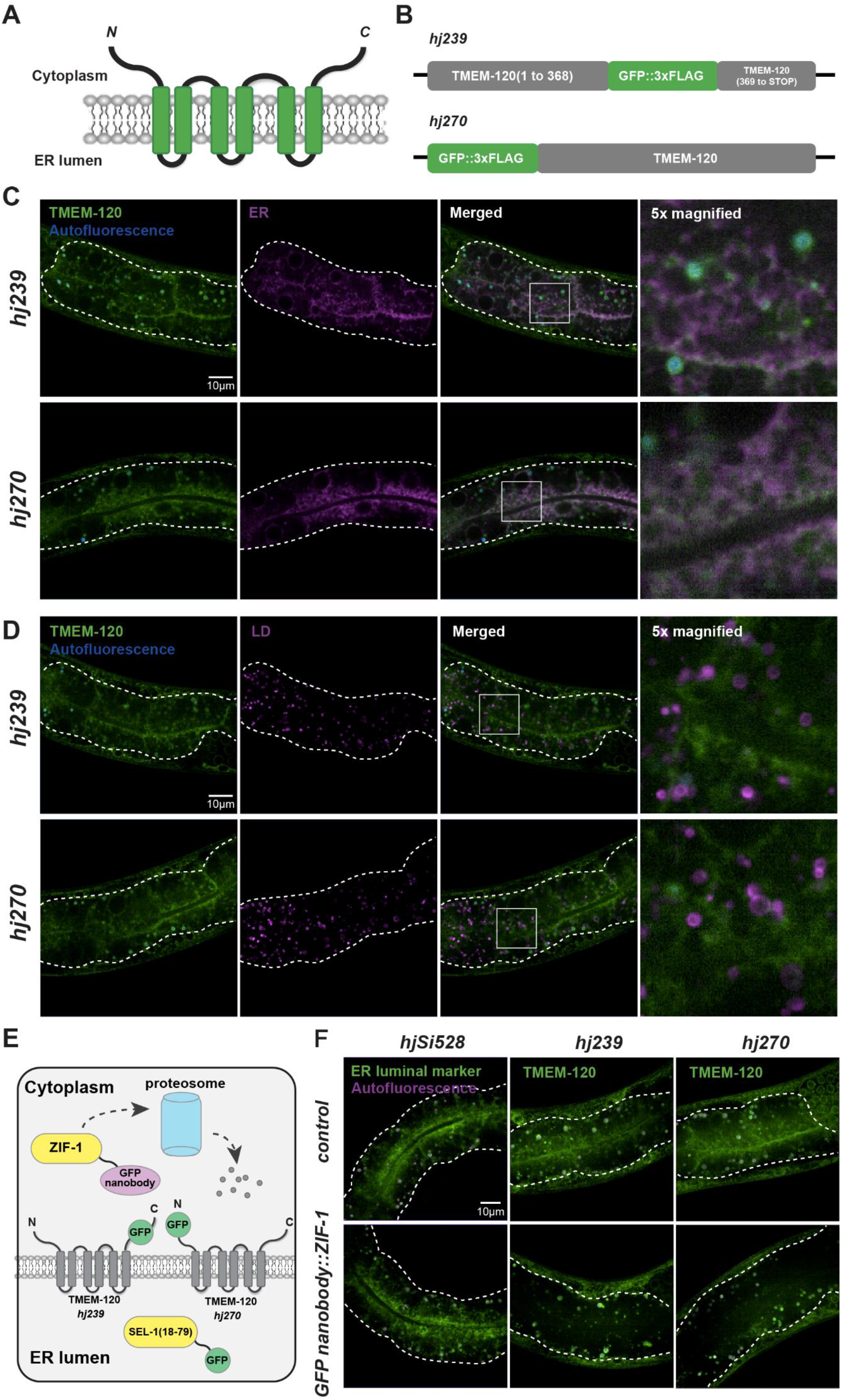
TMEM-120 is an ER resident protein. (A) The topology of TMEM-120 based on its homology with TMEM120A. (B) Schematic representation of TMEM-120 GFP fusion proteins. (C) Visualization of TMEM-120 GFP fusion proteins and an intestinal-specific luminal ER marker SEL-1(18-79)::mCherry::HDEL (*hjSi158*) in L4 worms. Representative images of a single focal plane of the first and second intestinal segment are shown. For each image, the intestine is enclosed by dashed lines. The boxed region in the merged image is 5x magnified and shown as a separate panel. (D) As in (C), but with an LD marker DHS-3::mRuby (*hj200*). (E) Schematic diagram on the GFP nanobody::ZIF-1 mediated degradation of cytoplasmic GFP fusion proteins. GFP targeted to the ER lumen is protected from degradation. (F) Visualization of an intestinal-specific luminal ER marker SEL-1(18-79)::GFP::HDEL (*hjSi528*) and TMEM-120 GFP fusion proteins in L4 worms in the absence (control) or presence of GFP nanobody::ZIF-1 (*hjSi524*). Each representative image is a projection of 5μm z stack with the second intestinal segment in the center. The intestine is enclosed by dashed lines. GFP signals in the hypodermis (regions outside the dashed lines) were unaffected by GFP nanobody::ZIF-1.

Next, we determined the localization of TMEM-120(G195E) by inserting the GFP coding sequence into the 5’ end of *tmem-120*(*hj49*). We found that GFP::TMEM-120(G195E) was expressed at a comparable level as the wild type protein and localized correctly to the ER (Fig. S3C). Therefore, we conclude that the G195E mutation most likely attenuates TMEM-120 function as predicted from the TMEM120A structure.

Finally, we experimentally verified the location of the N- and C-terminal tails of TMEM-120. To this end, we took advantage of the anti-GFP nanobody (vhhGFP4) directed protein degradation system (Wang et al., 2017). By expressing a GFP nanobody::ZIF-1 fusion protein in the intestinal cytoplasm, GFP fusion proteins with the GFP moiety exposed in the cytoplasm are subject to degradation (Fig. 3E). However, if the GFP moiety is in the ER lumen, the fusion protein is protected. Indeed, we detected fluorescence signals from a luminal ER GFP marker (*hjSi528*), even when GFP nanobody::ZIF-1 was co-expressed in the same worm (Fig. 3F). In contrast, the two versions of GFP fusion protein with TMEM-120 were subject to degradation, when GFP nanobody::ZIF-1 was co-expressed in the intestine (Fig. 3F). As a control, hypodermal GFP fusions with TMEM-120 remained detectable (Fig. 3F), indicating that anti-GFP nanobody directed, post-translational degradation was tissue-restricted as designed. Taken together, our results firmly suggest that TMEM-120 is an ER resident protein, with its N- and C-termini facing the cytoplasm.

### TMEM-120 regulates LD expansion cell autonomously

We generated an additional *tmem-120* deletion allele, *hj281*, using CRISPR mediated genome editing and Cre-LoxP mediated germline excision. Similar to the *hj50* allele, the *hj281* allele conferred *tmem-120* loss of function phenotypes and suppressed LD expansion in *daf-22* mutant worms (Fig. S3D-E). Re-expression of *tmem-120* in the intestine alone from a single-copy transgene (*hjSi557*) was sufficient to support LD expansion in *daf-22(-); tmem-120(hj281)* worms (Fig. S3D-E). Therefore, we conclude that TMEM-120 acts cell autonomously to promote LD expansion in *C. elegans*.

### TMEM120 promotes adipogenesis in mammalian cells

The expression level of TMEM120A and TMEM120B is induced during the differentiation of 3T3-L1 pre-adipocytes into adipocytes (Batrakou et al., 2015). In addition, knockdown of TMEM120A and TMEM120B impedes the differentiation of 3T3-L1 pre-adipocytes (Batrakou et al., 2015). To extend these observations, we used an alternative model of adipogenesis: the murine OP9 pre-adipocytes (Wolins et al., 2006). In the presence of insulin and oleic acid, the OP9 pre-adipocytes could differentiate into adipocytes in 3 days, which was accompanied by the induction of adipocyte markers such as Glut4 and adiponectin (Fig. S4A-B). Over the same time course, the expression level of TMEM120A and TMEM120B was significantly increased (Fig. 4A-B). Next, we stably knocked down the expression of TMEM120A and/or TMEM120B in OP9 cells by CRISPRi (Gilbert et al., 2013) (Fig. S4C-D). Wild type and knockdown cells were induced to differentiate and mature adipocytes were recognized by the appearance of a single, dominant LD (>15μm), as visualized by Oil Red O staining (Fig. 4C-D). Consistent with previous observations in 3T3-L1 cells, knockdown of TMEM120A and/or TMEM120B significantly reduced the ability of OP9 cells to differentiate into adipocytes (Fig. 4C-D). Our results suggest that the two mammalian TMEM120 paralogs are required for adipogenesis in vitro.

**Figure 4.**
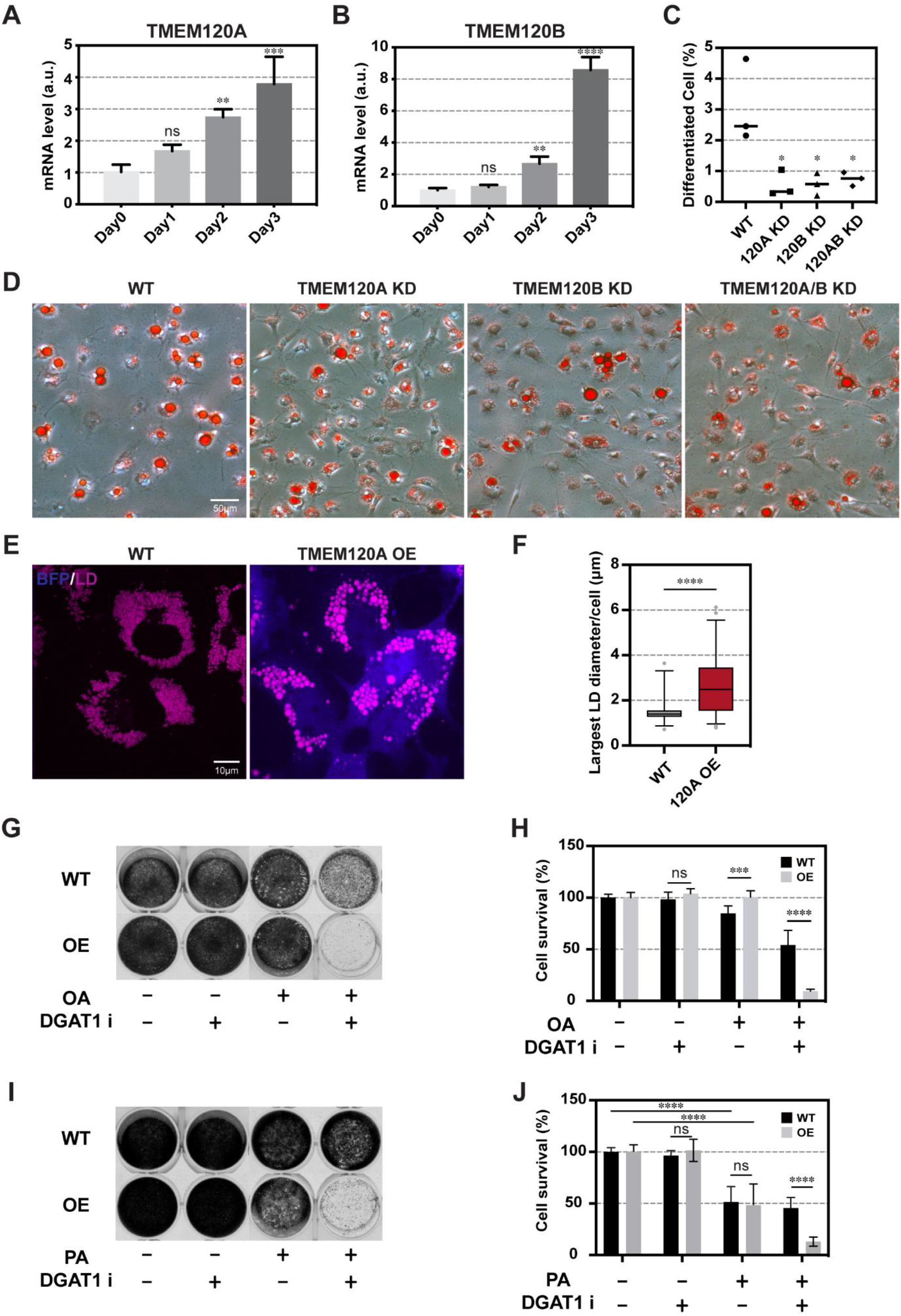
TMEM120A promotes adipogenesis and LD expansion in mammalian cells. (A-B) The expression level of TMEM120A and TMEM120B during OP9 pre-adipocyte differentiation measured by real time PCR. Mean + SD from three independent samples is shown. (C) Quantification of mature OP9 adipocytes (at least one LD >15μm / cell). Data summarized from three independent experiments. Each data point represents the percentage of mature adipocytes in one experiment. Total number of cells analyzed: WT, 4988 cells; TMEM120A KD, 5216 cells; TMEM120B KD, 6107 cells; TMEM120A+TMEM120B KD: 3898 cells. Horizontal line represents the mean. (D) Visualization of LDs in differentiated WT, TMEM120A KD, TMEM120B KD, and TMEM120A/B KD OP9 cells by Oil Red O staining. (E) Visualization of LDs in oleic acid treated wild type (WT) and TMEM120A overexpressing (OE) COS7 cells with FAS lipid droplet dye. Three independent experiments were performed. Each representative image is a projection of 5.5μm (WT) or 6.5μm (OE) z stack. (F) Quantification of the largest LD diameter of oleic acid treated WT and TMEM120A OE COS7 cells. Three independent experiments were performed. Total number of cells analyzed: WT: 183 cells. OE: 239 cells. (G) Assessment of cell survival by crystal violet staining after 20 hours of oleic acid (OA) and DGAT1 inhibitor (DGAT1i, A922500) treatment. Solvent control: ethanol (oleic acid) and DMSO (DGAT1i). (H) Quantification of cell survival, based on extracted crystal violet from (G). Data summarized from 9 independent wells, performed on 3 separate days, for each cell line. Mean + SD is shown. (I) As in (G), but with palmitic acid (PA). (J), As in (H), but with cells treated with palmitic acid (PA). Statistical analysis: (A-B) One-way ANOVA followed by Dunnett’s multiple comparisons test; (C and F) unpaired t test; (H-J) multiple t-test. ns, not significant; *p<0.05; **p<0.01; ***p<0.001; ****p<0.0001.

### TMEM120 promotes LD expansion in mammalian cells

Next, we tested if TMEM120A was sufficient to promote LD expansion in mammalian COS7 cells. To this end, we generated COS7 cells that stably over-expressed TMEM120A using the Sleeping Beauty Transposon system (Kowarz et al., 2015). When oleic acid was added to induce LD expansion, we observed significantly larger LDs in cells that over-expressed TMEM120A in comparison to parental cells (Fig. 4E-F). Therefore, TMEM120A is sufficient to promote LD expansion, plausibly by elevating the amount of fatty acids that are available for the synthesis of TAG, which is stored in LDs. It should be noted that excess fatty acids that cannot be incorporated into TAG, are toxic to cells. As a result, cell death ensues (Listenberger et al., 2003). Accordingly, COS7 cells that over-expressed TMEM120A were more sensitive than parental cells to inhibitors of diacylglycerol O-acyltransferase 1 (DGAT1) in the presence of exogenous fatty acids (Fig. 4G-J, S4E-F). Almost all cells that over-expressed TMEM120A died when oleic or palmitic acid was applied simultaneously with the DGAT1 inhibitor A922500. Interestingly, the dose-sensitive response to DGAT1 inhibitors was not observed for a diacylglycerol O-acyltransferase 2 (DGAT2) inhibitor (Fig. S4G). Based on the current model of LD biogenesis and expansion (Cao and Mak, 2020; Olzmann and Carvalho, 2019), DGAT1 in the ER acts early to synthesize TAG that supports the emergence of LDs from the ER while LD-localized DGAT2 contributes more significantly to the expansion of mature LDs. Our results on the differential sensitivity of TMEM120A over-expressing cells to DGAT1 inhibitors are consistent with the notion that TMEM120A acts in the ER to promote the incorporation of fatty acids into TAG, upstream of DGAT1.

### Concluding remarks

In this paper, we combined genetic, imaging and pharmacological approaches to demonstrate an evolutionarily conserved function of TMEM-120/TMEM120A in promoting TAG synthesis and LD expansion. Our examination of TMEM-120 at the endogenous level in live worms strongly suggests the ER as its primary site of action. This is consistent with the notion that the ensemble of TAG synthesis proteins can all be found in the ER, which conceivably enables the transfer of biosynthetic intermediates within the membrane. Although we do not yet know if TMEM-120/TMEM120A acts specifically with one or more enzymes in the TAG synthesis pathway, the ability of purified TMEM120A to bind CoASH is intriguing (Niu et al., 2021; Rong et al., 2021; Xue et al., 2021). This is because fatty acids are converted to fatty acyl-CoA by acyl-CoA synthetases, prior to their addition to the glycerol backbone by a series of acyltransferases (Coleman and Lee, 2004; Yen et al., 2008). We note that TMEM-120 mutant worms remain capable of TAG synthesis, albeit at a reduced level (Fig. 1E). It is plausible that TMEM-120/TMEM120A can trap CoASH or acyl-CoA in the ER to enhance the efficiency of TAG synthesis. Such hypothesis can be tested with purified proteins and substrates in vitro. In conclusion, our functional studies in *C. elegans* and mammalian cells support the conclusion from structural studies that TMEM120A and its orthologs are unlikely to be mechanosensory channels.

## Materials and Methods

### Strains and transgenes

The wide type *C. elegans* strain was Bristol N2. All experimental animals were maintained at 20°C. The following strains and alleles were used: LG I, EG6701 (*ttTi4348*); LG II, *daf-22* (*ok693*); LG III, *tmem*-120(*h49*), *tmem-120*(*hj50*); LG IV, EG6703 (*cxTi10816*).

The following transgenes or CRISPR-generated alleles were used:

*hjSi158 [vha-6p::sel-1(a.a.1-79)::mCherry::HDEL::let-858 3’UTR] I*
*hjSi524 [vha-6p::vhhgfp4::zif-1::let-858 3’UTR] I*
*hjSi528 [vha-6p::sel-1(a.a.1-79)::GFP::HDEL::tbb-2 3’UTR] IV*
*hjSi557 [vha-6p::gfp::3xFLAG::tmem-120(codon modified) cDNA::dhs-28 3’-UTR] IV dhs-3 (hj200) [dhs-3::mRuby] I*
*tmem-120 (hj239) [tmem-120(a.a.1-368)::gfp::3xFLAGG::tmem-120(a.a.369 to stop)] III*
*tmem-120 (hj266) [tmem-120(a.a.1-368)(loxP in introns 3 and 4)::gfp::3xFLAG::tmem-120(a.a.369 to stop)] III*
*tmem-120 (hj270) [GFP::3xFLAG::tmem-120] III*
*tmem-120 (hj297) [GFP::3xFLAG::TMEM-120(G195E)] III*

The *tmem-120*(*hj281*) allele was generated by Cre-loxP mediated germline excision of *tmem-120*(*hj266*).

Extrachromosomal array strains:

*hjEx29[vha-6p::tmem-120 cDNA::sl2::gfp::let-858 3’ UTR]*
*hjEx30[vha-6p::mouse tmem120a cDNA::sl2::gfp::let-858 3’UTR]*

All strains were outcrossed with wild type N2 at least twice before further characterization.

### Genetic screen

The *tmem-120* alleles were isolated in a genetic screen for suppressors of the expanded LD phenotype of *daf-22*(*ok693*) mutants. Complementation tests indicated that *hj49* and *hj50* belonged to the same complementation group. Using a SNP-based mapping strategy with the Hawaiian *C. elegans* isolate CB4856 (Davis et al., 2005), we mapped *hj50* to LGIII. Mutant worms were then subjected to whole genome sequencing (Illumina). The molecular lesions of *hj49* and *hj50* in M01G5.3 were confirmed by Sanger sequencing. The *daf-22(ok693); tmem-120(hj50)* mutant could be rescued by a *vha-6p::tmem-120::sl2::gfp* transgene (Fig. 1F).

### Fluorescence Imaging of *C. elegans* and mammalian cells

Fluorescence imaging of live worms and mammalian cells was performed as described (Cao et al., 2019). In brief, fluorescence images of worms at indicated stages were acquired on a spinning disk confocal microscope (AxioObeserver Z1, Carl Zeiss) equipped with a piezo Z stage using a 63x, NA 1.4 (for visualizing TMEM-120 localization) or 100x, NA 1.46 (for quantification of LD diameter) oil Alpha-Plan-Apochromat objective on a Neo sCMOS camera (Andor) controlled by the iQ3 software (Andor). For GFP, a 488nm laser was used for excitation and signals were collected with a 500-550nm emission filter. For mRuby or mCherry, a 561nm laser was used for excitation and signals were collected with a 580.5-653.5nm emission filter. For autofluorescence from lysosome-related organelles, a 488nm laser was used for excitation and signals were collected with a 580.5-653.5nm emission filter. For BFP, a 405 nm laser was used for excitation, and signals were collected with a 437nm emission filter. For FAS LD dye, a 405nm laser was used for excitation and signals were collected with a 617nm emission filter. Optical sections, as specified, were taken at 0.5μm intervals and z stacks of 8μm-10μm were exported from iQ3 to Imaris 8 (Bitplane) for processing.

### Live imaging by Stimulated Raman Scattering

Stimulated Raman Scattering (SRS) for measuring endogenous neutral lipid in live worms was carried out as described (Li et al., 2015). Live animals at specific stages were mounted on 8% agarose pad in 1xPBS buffer with 0.2mM levamisole. For visualizing LDs, the focal plane for the center of the first and second intestinal segments in young adult animals was determined with a 40x water immersion objective (UAPO40XW3/340, 1.15 NA, Olympus). To measure neutral lipid content level in whole worms, the focal plane with maximal SRS intensity was determined with a 20x air objective (Plan-Apochromat, 0.8 NA, Zeiss). The C-H bound was detected at 2863.5 cm^−1^. The quantification of SRS signal was done following a published protocol (Ramachandran et al., 2015). SRS of live worms for measuring fatty acid absorption and lipolysis were carried out as described with a 40x water immersion objective (UAPO40XW3/340, 1.15 NA, Olympus) (Li et al., 2019). Saturated bacterial cultures of OP50 were mixed with 4mM deuterium labeled PA-d_31_ or OA-d_34_ (Sigma) and then seeded onto NGM plates. To measure fatty acid uptake, populations of young adult worms (before egg-laying started), raised in the absence of deuterium labeled fatty acid were transferred to plates with PA-d_31_ or OA-d_34_. To measure lipolysis, populations of L1 larvae were raised on plates with PA-d_31_ or OA-d_34_, and transferred to OP50 seeded plates without deuterium labeled fatty acid when they were young adults. Animals were imaged by SRS at 5 to 6-hour intervals for 28 to 30 hours. The C-D bond vibration was detected at 2116.8 cm^−1^. Images were imported into ImageJ and further processed for quantification in MATLAB_R2015a.

### Real-time PCR for *C. elegans* samples

For each experimental sample, around 400 worms were synchronized at the L1 larval stage. Worms were harvested at the L4 stage (43 hours after L1 for WT, and 46 hours after L1 for *tmem-120*(*hj50*)), and total RNA extracted with a Direct-zol™ RNA MiniPrep kit (Zymo). 200ng of total RNA was reverse transcribed with a Transcriptor cDNA synthesis kit (Roche). The real time PCR was carried out on a Roche LightCycler system with SYBR Green Master Mix (Roche). For each strain, technical triplicates were performed for each biological sample. The Delta-delta CT method was used for analyzing the raw CT values.

The following primers were used for RT-qPCR:

For *tmem-120*:

Forward: 5’-TGAGACAAGCCCAACAATCA-3’
Reverse: 5’-TGGAGCCCAAAATCAAATTC-3’

For internal standard *rpl-32*:

Forward: 5’-AGGGAATTGATAACCGTGTCCGCA-3’
Reverse: 5’-TGTAGGACTGCATGAGGAGCATGT-3’

### Real-time PCR of mammalian samples

Total RNA was extracted from mammalian cells using Direct-zol™ RNA MiniPrep kit (Zymo) following the manufacturer’s protocol. 500ng of total RNA was reverse transcribed using First Strand cDNA Synthesis Kit (Sigma Aldrich) following the manufacturer ‘s protocol. The real time PCR was performed on a LightCycler480 system (Roche) using SYBR Green Master Mix (Roche) following the manufacturer’s protocol. Data was obtained with 3 biological samples of each cell line, tested in technical triplicates. The Delta-delta CT method was used for analyzing the raw CT values.

The following primers were used:

For TMEM120A:

Forward: 5’-AGGGCTTTCAGTCTTGGATG-3’
Reverse: 5’-AAATTGCCGAGGAAGAGGAG-3’

For TMEM120B:

Forward: 5’-TCAGAGCTGCGTTCAGTTTC-3’
Reverse: 5’-ACAGAAGAGGAAAGGCAGGAG-3’

For Glut4:

Forward: 5’-GTAACTTCATTGTCGGCATGG-3’
Reverse: 5’-CTCTGGTTTCAGGCACTTTTAG-3’

For Adiponectin:

Forward: 5’-CCTGGCCACTTTCTCCTC-3’
Reverse: 5’-GTGGAGGGACCAAAGCAG-3’

For 36B4 (internal control) (Zhang et al., 2016):

Forward: 5’-CTGAGTGATGTGCAGCTGAT-3’
Reverse: 5’-AGAAGGGGGAGATGTTCAG-3’

### Pharyngeal pumping rate

Videos for pharyngeal pumping rate measurement were obtained on an OLYMPUS SZX16 stereo microscope. One day before the imaging, L4 stage worms were transferred to a newly seeded NGM plate. The plates were left beside the microscope for acclimatization. For each experimental group, up to 10 worms were prepared. A 2-minute video focusing on the pharynx of each worm was captured. Each video was trimmed to a 60 seconds clip and then played under 0.3X speed for counting the number of pharyngeal contractions visually.

### Lipid analysis

Lipid extraction was conducted using methyl-tert-butyl ether(MTBE) as described (Matyash et al., 2008; Witting et al., 2014), with modifications. For each experimental sample, around 2,000 worms were synchronized at L1 larvae stage. Worms were harvested at the L4 stage (43 hours after L1 for WT, and 46 hours after L1 for *tmem-120*(*hj50*)), washed with detergent free PBS for at least three times and transferred into organic solvent resistant Eppendorf tubes. 250μl methanol (precooled to −20°C) (RDH, for HPLC) was added and samples were frozen in liquid nitrogen and stored at −80°C. For extraction, samples were thawed on ice, and 875μl MTBE (VWR, for HPLC) was added. Worms were lysed with ice cold ultrasonic bath with an interval of 2 mins on and 30 seconds off. Phase separation was induced by the addition of 210μl water with further sonication for 15 mins. After centrifugation at 16,100xG at 4°C for 15 mins, the upper organic phase was collected into a glass vial. 325μl MTBE was added to the lower phase and centrifuged at 17,000xG at 4°C for 15 mins for re-extraction of lipids. Upper organic phase was collected after centrifugation and combined with those previously collected. Extracts were dried under a stream of nitrogen at room temperature, re-dissolved in 200μl acetonitrile (RCI Labscan, for HPLC)/ isopropanol (RNH)/ water(65/30/5, v/v/v) and stored at −80 °C.

Lipid analysis was performed as previously described (Zeng et al., 2020). A 100μl aliquot was analyzed using the Bruker Elute UPLC system with 2 technical injections per sample. The mass spectrometry data was analyzed with MetaboScape version 5.0, annotated with spectral libraries MSDIAL-Tandem Mass Spectral Atlas-VS68-pos and MSDIAL- Tandem Mass 736 Spectral Atlas-VS68-neg. The intensity was normalized with probabilistic quotient normalization method (Dieterle et al., 2006). Signals from all TAG species were summed.

### Cell culture

OP9 mouse stromal cells (ATCC-CRL-2749) were maintained in α-MEM (Life Technologies) with 20% FBS (Life Technologies) and 1% antibiotic-antimycotic (Life Technologies). COS7 Cells (ATCC-CRL-1651) were maintained in DMEM (Life Technologies) with 10% FBS (Life Technologies) and 1% antibiotic-antimycotic (Life Technologies). All the cell lines were incubated in 37°C humidified incubator with 5% CO_2_.

### Generation of TMEM120A overexpressing COS7 cells

The Sleeping beauty transposon system was used to generate COS7 cells that overexpressed human TMEM120A. Cells were co-transfected with sleeping beauty transposon plasmid pSBi-Hyg-BFP (hTMEM120A cDNA) and sleeping beauty transposase plasmid pCMV (CAT) T7-SB100 with Lipofectamine2000 (Life Technologies). Three days after transfection, cells were maintained in selection medium (400μg/mL hygromycin in DMEM growth medium) for at least seven days. Cells that survived drug selection were sorted using Aria III system (Becton Dickinson) and sub-divided into ‘Low’, ‘Medium’, and ‘High’ populations based on BFP fluorescence intensity (a surrogate of TMEM120A expression). The ‘High’ cell population was used in subsequent experiments.

### Generation of TMEM120A and TMEM120B knockdown OP9 cells

OP9 cells were co-transfected with the sleeping beauty transposase plasmid pCMV (CAT) T7-SB100 and the transposon plasmid (Krab::dCas9::BFP::TMEM120A sgRNA or TMEM120B sgRNA). The sgRNA sequences were selected from a published database (Horlbeck et al., 2016). The transfected cells were maintained in selection medium (400μg/mL hygromycin in α-MEM growth medium) for at least seven days. Cells that survived drug selection were sorted into ‘Low’, ‘Medium’, and ‘High’ populations based on the BFP fluorescence intensity (a surrogate of dCas9 and sgRNA expression). The ‘Medium’ cell populations were used for subsequent experiments.

For the TMEM120A+TMEM120B double knockdown cells, the sleeping beauty transposase plasmid and the transposon plasmid (mRuby::TMEM120B sgRNA) were transfected into TMEM120A knockdown cells. Drug selection and cell sorting were performed as described above except that puromycin was used instead of hygromycin. The ‘High’ cell population was used for subsequent experiments. The knockdown efficiency of all stable cell lines was determined by real time PCR.

### Fatty acid supplementation

Fatty acids supplementation was performed according to a published method (Cao et al., 2019; Peng et al., 2011). In brief, the cell culture medium was pre-heated to 60°C for 5 mins prior to the addition of fatty acids (400μM oleic acid or palmitic acid). The medium was then equilibrated to 37°C before use. To induce LD expansion, cells were incubated with 400μM FAs for 20 hours. To visualize LDs, the cells were stained with 10μM FAS (Wang et al., 2016) for 15 mins prior to imaging.

### DGAT inhibitor and fatty acids treatment

The cell culture medium was pre-heated to 60°C for 5 mins prior to the addition of fatty acids (400μM oleic acid or 400μM palmitic acid or ethanol control), without (DMSO control) or with DGAT inhibitors as specified (A922500, PF04620110 or PF 06424439 (R&D system)). The medium was equilibrated to 37°C before use. The cells were treated with 20 hours (oleic acid) or 60 hours (palmitic acid) before they were stained with crystal violet. Quantification of crystal violet was performed according to a published protocol (Feoktistova et al., 2016).

## Author Contributions

Y.L. was responsible for Figs. 1, 2, 3, S1 (except S1F), S2 and S3. S.H. was responsible for Figs. 4 and S4. X.L. and J.Q. were responsible for developing the SRS system and Figs. 1A-B, 2 and S2. X.Y. was responsible for Fig. S1F and the generation of *tmem-120*(*hj266*), *tmem-120*(*hj281*) and *hjSi557*. N.X. conducted genetic screen, genetic mapping and molecular cloning of *C. elegans* mutants. H.Y.M. supervised the project and wrote the paper.

## Acknowledgements

We thank Yifei Qiu for preliminary data, Zhenfeng Liu for discussion, Pui Shuen Wong at the HKUST Bioscience Central Research Facility for lipidomic analysis, the Molecular Biology core at the Stowers Institute for Medical Research for whole-genome sequencing. Some strains were provided by the CGC, which is funded by NIH Office of Research Infrastructure Programs (P40 OD010440). This work was supported by RGC GRF 662013 to HYM.

---

**Figure S1.**
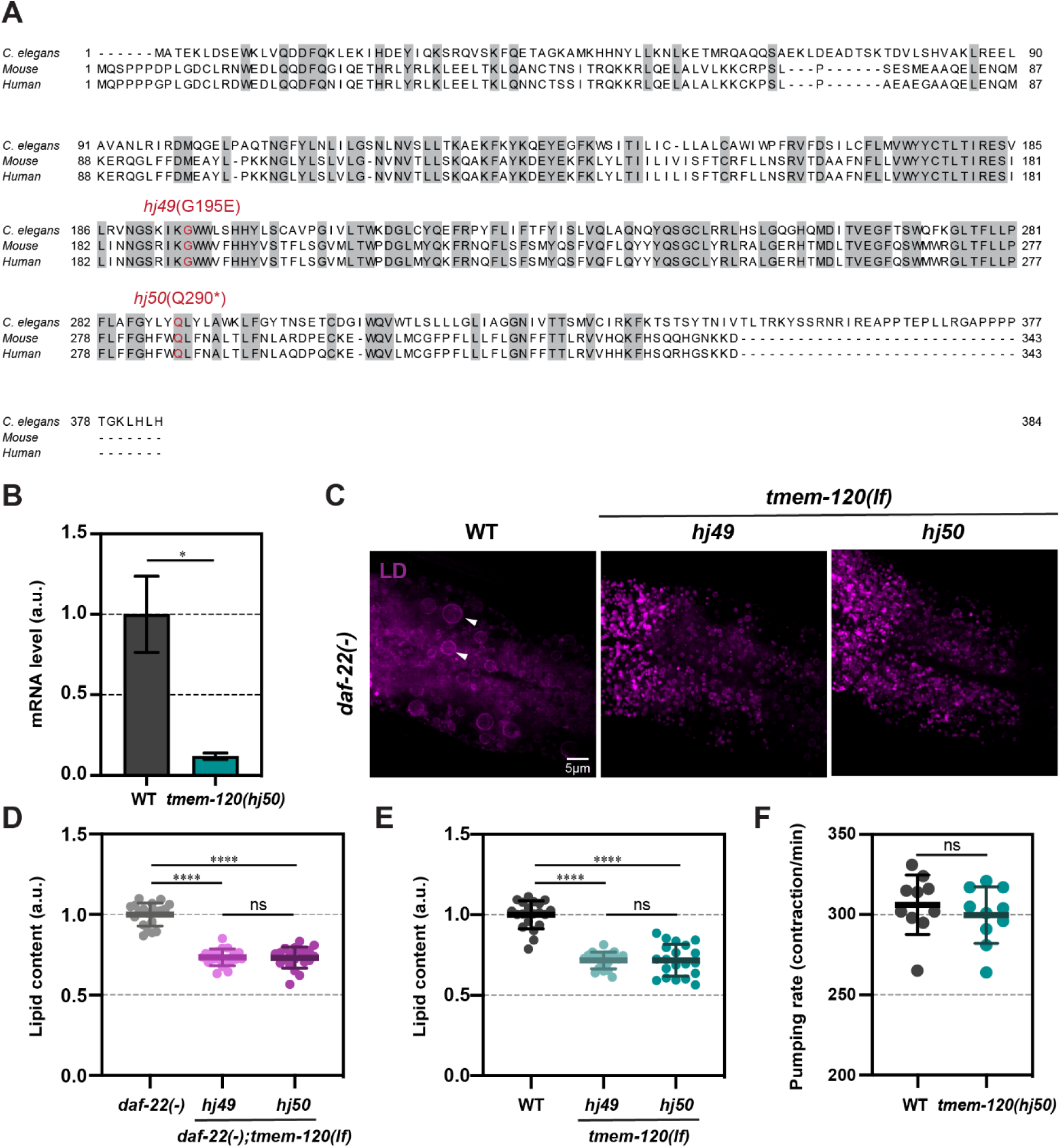
TMEM-120 promotes LD expansion. (A) Sequence alignment of *C. elegans* TMEM-120, mouse TMEM120A and human TMEM120A. Identical residues are shaded in grey. The mutated residues encoded by *tmem-120*(*hj49*) and *tmem-120*(*hj50*) are labeled red. (B) Expression level of *tmem-120* in wild type (WT) and *tmem-120(hj50)* L4 stage worms measured by real time PCR. Two independent samples for each strain. The mean value of WT is set as 1 for comparison. (C) Visualization of LDs in *daf-22(-), daf-22(-); tmem-120(hj49)* and *daf-22(-); tmem-120(hj50)* larval L4 stage worms using DHS-3::mRuby (*hj200*) as a LD marker. Each representative image is a projection of 7.5μm z stack with the second intestinal segment in the center. Enlarged LDs are indicated by white arrowheads. (D) Label free quantification of neutral lipid content in *daf-22*(-) (n=20), *daf-22*(-); *tmem-120*(*hj49*) (n=19) and *daf-22(-); tmem-120*(*hj50*) (n=20) L4 stage worms by SRS. Each data point represents the neutral lipid content of an individual worm. (E) As in (D), but with WT (n=20), *tmem-120*(*hj49*) (n=19) and *tmem-120*(*hj50*) (n=20) L4 stage worms. (F) Pharyngeal pumping rate of WT (n=10) and *tmem-120*(*hj50*) (n=10) 1-day old adult worms. Each data point represents the contraction rate of an individual worm. For all graphs, bars or horizontal lines represent mean ± SD. Statistical analysis: (B and F) unpaired t test; (D and E) one-way ANOVA followed by Tukey’s multiple comparisons test. a.u., arbitrary unit. ns, not significant; *p < 0.05; ****p < 0.0001.

**Figure S2.**
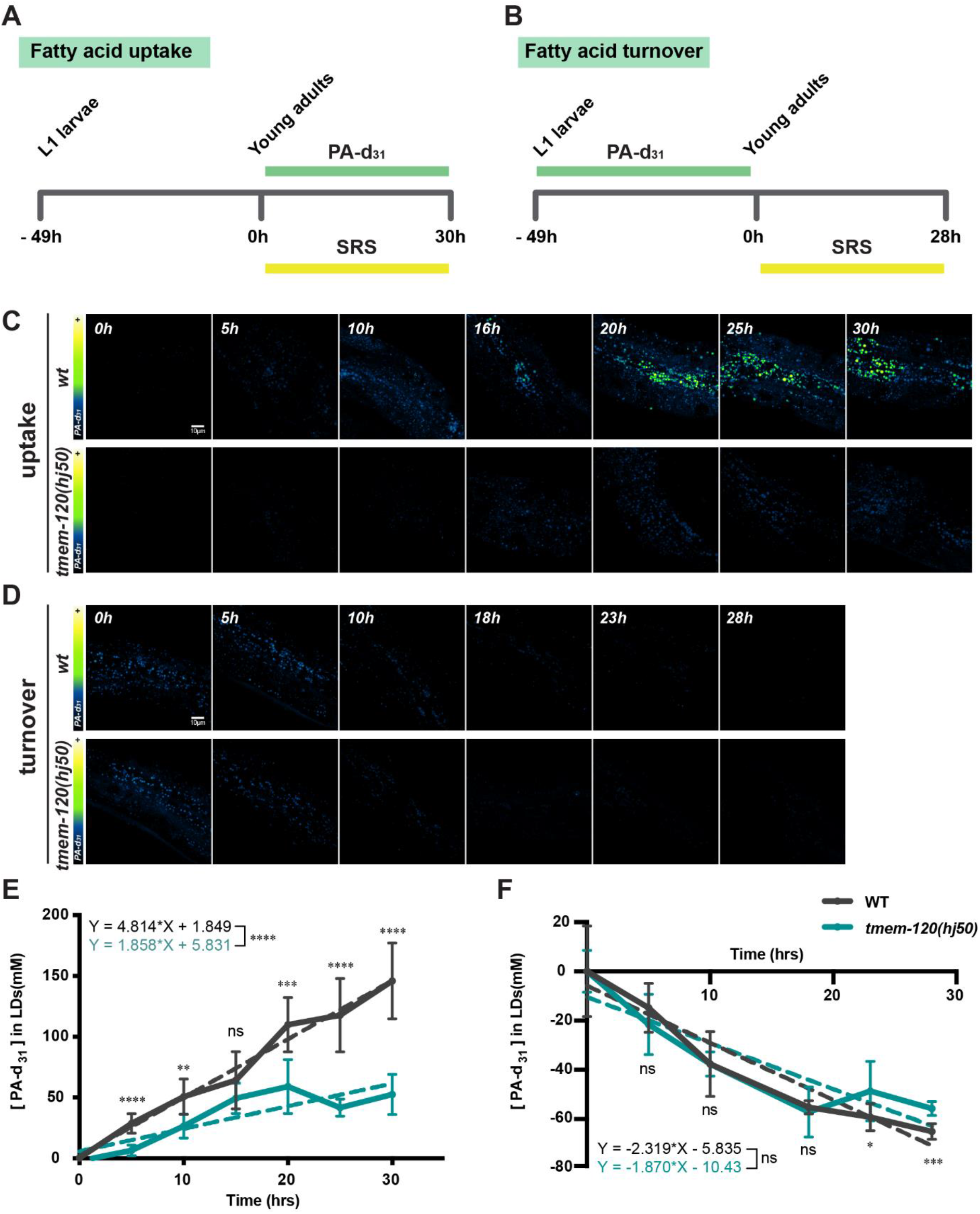
TMEM-120 promotes the incorporation of fatty acids into TAG. (A-B) The experimental design for monitoring deuterated palmitic acid-d_31_ (PA-d_31_) uptake (A) or turnover (B). (C-D) Visualization of PA-d_31_ incorporation (C) or turnover (D) by SRS in wild type (wt) and *tmem-120*(*hj50*) worms. Representative images of a layer with the strongest SRS signal in the first and second intestinal segments are shown. (E) Quantification of PA-d_31_ uptake in WT and *tmem-120*(*hj50*) worms shown in (C). n = 5 to 9 for each group at each time point. Straight dashed lines and equations were generated based on linear regression analysis of each group. (F) Quantification of PA-d_31_ turnover in WT and *tmem-120*(*hj50*) worms shown in (D). n = 4 to 9 for each group at each time point. Statistical analysis: unpaired t test (for each time point). ns, not significant; *p < 0.05; **p < 0.01; ***p < 0.001; ****p < 0.0001.

**Figure S3.**
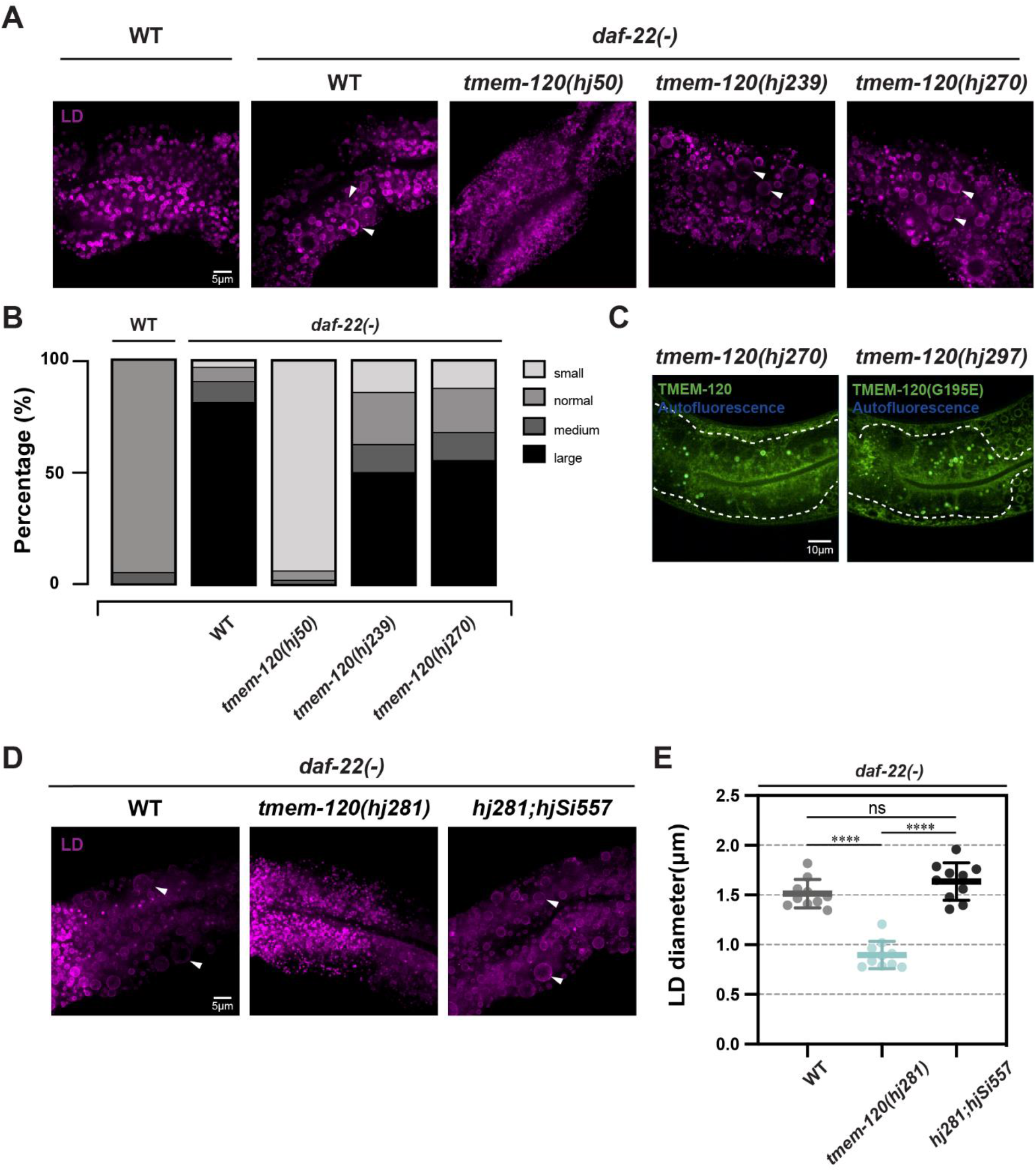
Functional characterization of TMEM-120 GFP fusion proteins. (A) Visualization of LDs in wild type (WT), *daf-22(-), daf-22(-); tmem-120(hj50), daf-22(-); tmem-120(hj239)* and *daf-22(-); tmem-120(hj270)* L4 stage worms, using DHS-3::mRuby (*hj200*) as a LD marker. Each representative image is a projection of 7.5μm z stack with the second intestinal segment in the center. Enlarged LDs are indicated by white arrowheads. (B) Quantification of percentage of WT (n=57), *daf-22*(-) (n=64), *daf-22(-); tmem-120(hj50)* (n=48), *daf-22(-); tmem-120(hj239)* (n=56) and *daf-22(-); tmem-120(hj270)* (n=56) 5-day old adult worms with small (D_L_ < 1.0μm), normal (1.0μm < D_L_ < 3μm), medium (3μm < D_L_ < 5μm) and large (5μm < D_L_) LDs. D_L_, diameter of the largest LD in the second intestinal segment of an individual worm. Data combined from 3 independent groups of worms (n = 14 to 22 for each group) for each strain. (C) Visualization of GFP::TMEM-120 (*hj270*) and GFP::TMEM-120(G195E) (*hj297*) in L4 stage worms. Representative images of a single focal plane of the first and second intestinal segments are shown. The intestine is enclosed by dashed lines. (D) Visualization of LDs in *daf-22*(-) and *daf-22(-); tmem-120(hj281)* and *daf-22(-); tmem-120(hj281); hjSi557[vha-6p::gfp::tmem-120]* L4 stage worms, using DHS-3::mRuby (*hj200*) as a LD marker. *tmem-120*(*hj281*) was generated by Cre-loxP based excision of *tmem-120*(*hj239*). Each representative image is a projection of 7.5μm z stack with the second intestinal segment in the center. Enlarged LDs are indicated by white arrowheads. (E) Quantification of LD size of worms from (D). Each data point represents average LD diameter of an individual worm. Total number of LDs quantified: *daf-22*(-) = 1276, *daf-22(-); tmem-120*(*hj281*) = 1216 and *daf-22(-); tmem-120(hj281); hjSi557* = 1279. Horizontal bars represent mean ± SD. Statistical analysis: one-way ANOVA followed by Tukey’s multiple comparisons test. ns, not significant; ****p < 0.0001.

**Figure S4.**
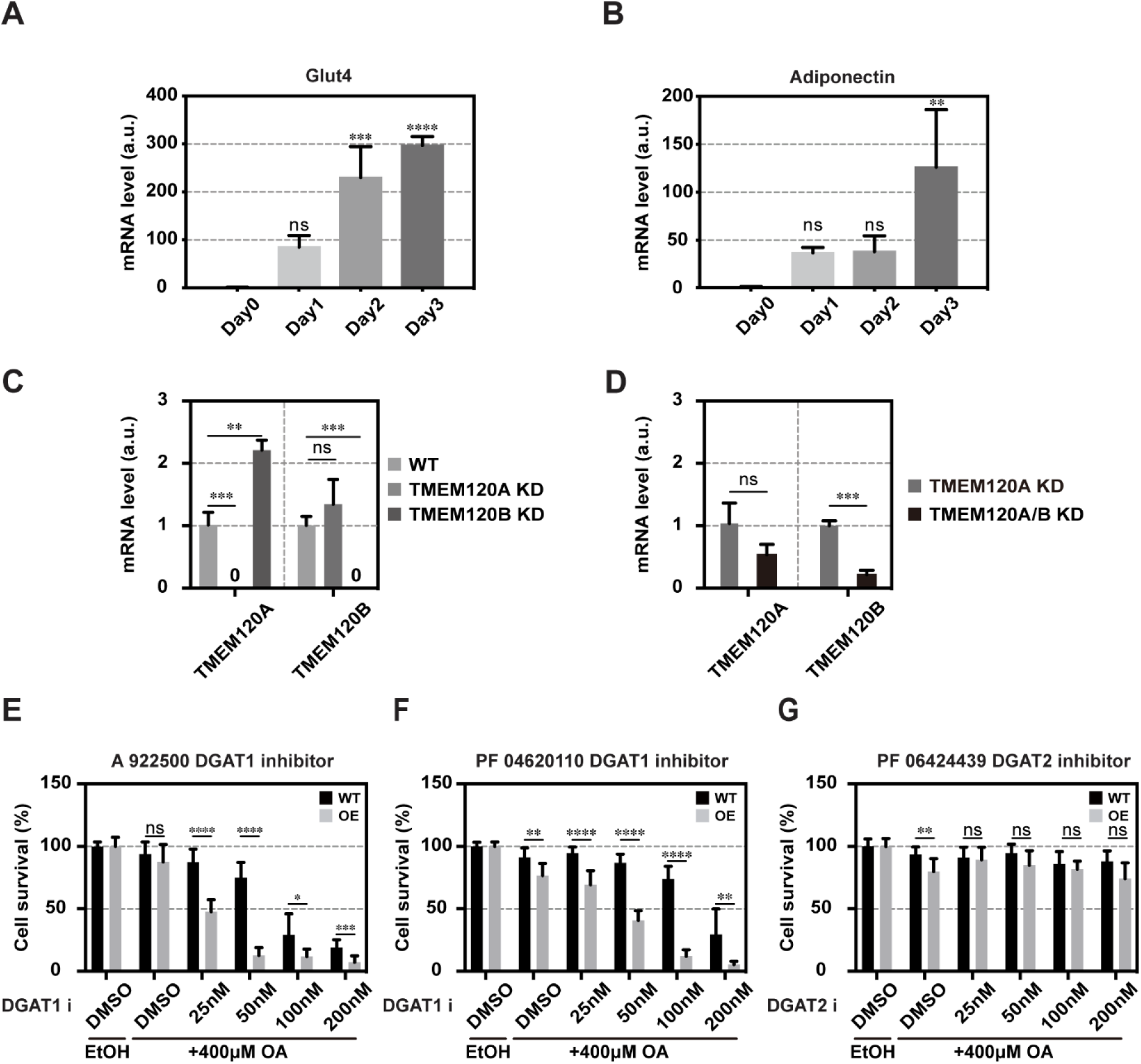
TMEM120A acts upstream of DGAT1 in mammalian cells. (A-B) The expression level of mature adipocyte markers Glut4 and Adiponectin during OP9 pre-adipocyte differentiation measured by real time PCR. Mean + SD from three independent samples is shown. The mean value on Day 0 is set as 1. a.u., arbitrary unit. (C) The expression level of TMEM120A and TMEM120B in wild type (WT), TMEM120A KD, TMEM120B KD cells, measured by real time PCR. Mean + SD from three independent samples of each cell line is shown. The mean value of WT cells is set as 1. (D) The expression level of TMEM120A and TMEM120B in TMEM120A KD (parental to the double KD cells) and TMEM120A+TMEM120B KD cells, measured by real time PCR. Mean + SD from three independent samples of each cell line is shown. The mean value of TMEM120A KD cells is set as 1. (E) Quantification of cell survival for WT and TMEM120A overexpressing (OE) COS7 cells treated with oleic acid (OA) and increasing concentration of DGAT1 inhibitor (A922500), based on crystal violet staining. Solvent control: ethanol (oleic acid) and DMSO (DGAT1i). Data summarized from 9 independent wells, performed on 3 separate days, for each cell line. Mean + SD is shown. (F) As in (E), but with DGAT1 inhibitor (PF04620110). (G) As in (E), but with DGAT2 inhibitor (PF06424439). Statistical analysis: (A-B) One-way ANOVA followed by Dunnett’s multiple comparisons test. (C-D) unpaired t-test; (E-G) multiple t-test. ns, not significant; *p<0.05; **p<0.01; ***p<0.001; ****p<0.0001.

